# The Project MinE databrowser: bringing large-scale whole-genome sequencing in ALS to researchers and the public

**DOI:** 10.1101/377911

**Authors:** Project MinE ALS Sequencing Consortium

**Author notes:** A full list of Project MinE GWAS Consortium members appears at the end of the paper. Corresponding author: Jan H. Veldink, Department of Neurology and Neurosurgery, University Medical Centre Utrecht, Department of Neurology G03.228, P.O. Box 85500, 3508 GA Utrecht, The Netherlands. shared first. shared last.

## Abstract

Amyotrophic lateral sclerosis (ALS) is a rapidly progressive fatal neurodegenerative disease affecting 1 in 350 people. The aim of Project MinE is to elucidate the pathophysiology of ALS through whole-genome sequencing at least 15,000 ALS patients and 7,500 controls at 30X coverage. Here, we present the Project MinE data browser (databrowser.projectmine.com). a unique and intuitive one-stop, open-access server that provides detailed information on genetic variation analyzed in a new and still growing set of 4,366 ALS cases and 1,832 matched controls. Through its visual components and interactive design, the browser specifically aims to be a resource to those without a biostatistics background and allow clinicians and preclinical researchers to integrate Project MinE data into their own research. The browser allows users to query a transcript and immediately access a unique combination of detailed (meta)data, annotations and association statistics that would otherwise require analytic expertise and visits to scattered resources.

## Introduction

Amyotrophic lateral sclerosis (ALS) is a rapidly progressive fatal neurodegenerative disease affecting 1 in 350 people. While research over the past years has revealed an increasing number of genetic variants contributing to ALS risk, the bulk of heritability in ALS remains to be elucidated. In addition to known rare variants, there is evidence for a central role of low-frequency and rare genetic variation in ALS susceptibility (van Rheenen et al., 2016). Well-powered genetic studies enabled through large-scale collaboration are crucial for identify these variants and improving our understanding of ALS pathophysiology (Benyamin et al., 2017; Schijven et al., 2017).

Project MinE, an international collaboration, was initiated precisely with the challenge of sample aggregation in mind. The Project MinE ALS sequencing Consortium has set out to collect whole-genome sequencing (WGS) of 15,000 ALS patients and 7,500 controls(van Rheenen et al., 2018). Currently, the Project MinE initiative has sequenced 4,366 ALS patients and 1,832 age- and sex-matched controls. Project MinE is a largely crowd-funded initiative. As such, we are committed to sharing data and results with the scientific and healthcare communities, as well as the public more broadly. Data sharing within the genetics community facilitated large-scale genome-wide association studies and ignited initiatives such as the Gene Atlas, LDhub GWAShare Center, and MRbase, places where people can share, explore and analyze data with few restrictions (Canela-Xandri et al., 2017) (Zheng et al., 2017). In this same spirit, we aim to share raw sequence data, provide results from our analyses, and facilitate interpretation through integration with existing datasets to serve researchers and the public across disciplines.

We, therefore, created the Project MinE databrowser (databrowser.projectmine.com). We integrated multi-level association statistics, metadata, and public resources including gnomAD, GTEx and ClinVar in an intuitive and flexible framework (**Fig.1**) (Lek et al., 2016),(GTEx Consortium, 2013),(Landrum et al., 2016). These data are freely available through the browser for any research initiative. We aim for the data to serve several purposes, including providing a backbone for new gene discovery, serving as a costless replication dataset, and aiding clinical interpretation of individual ALS patient genomes or specific genetic variants.

**Figure 1.**
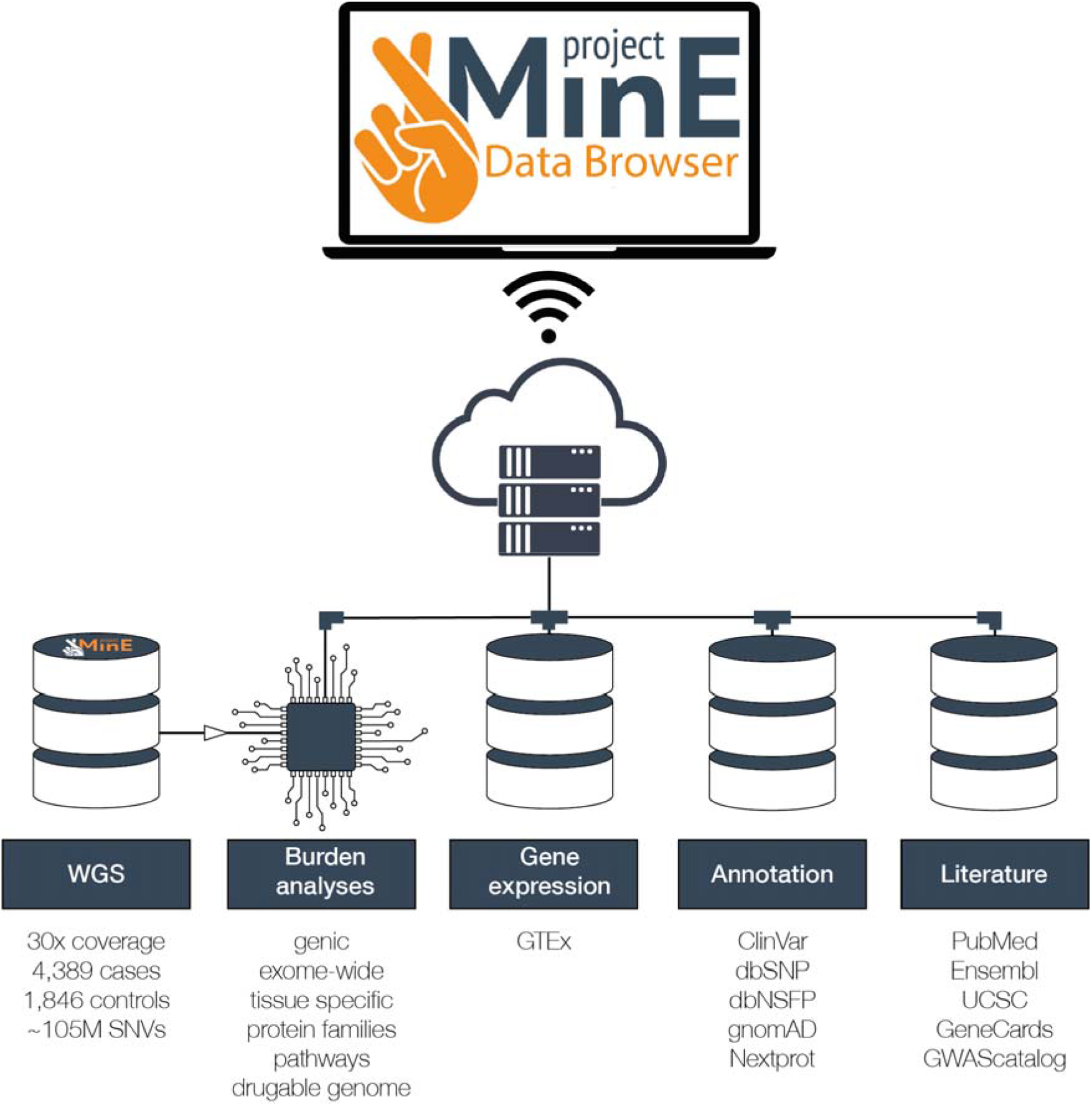
Schematic representation of the databrowser. Whole genomes generated by Project MinE are openly available for research and the public. The databrowser does not have a login requirement. It integrates multiple public resources and provides a wide range of robust statistical analyses.

## Materials and Methods

### Sample selection and WGS

The first batch of samples (1,935 cases and controls collected in the Netherlands) were sequenced on the Illumina HiSeq 2000 platform(van Rheenen et al., 2018). All remaining samples (4,644 cases and controls) were sequenced on the Illumina HiSeq X platform. All samples were sequenced to ~35X coverage with 100bp reads for the HiSeq 2000 and ~25X coverage with 150bp reads for the HiSeq X. Both sequencing sets used PCR-free library preparation. Samples were also genotyped on the Illumina 2.5M array. Sequencing data was then aligned to GRCh37 using the iSAAC Aligner, and variants called using the iSAAC variant caller; both the aligner and caller are standard to Illumina’s aligning and calling pipeline. Additional information regarding data merging, Sample- and variant level quality control can be found in the Supplementary Information.

### Association analyses

The main association analysis consists of several rare-variant burden analyses for an association with ALS risk. For quality control we have performed single variant association analysis using a mixed linear model, including a genetic relationship matrix and the first 20 PCs, as implemented in GCTA(Yang et al., 2011). We set genome-wide significance in single variant association analyses at p < 5 × 10^−9^ (Pulit et al., 2016), to account for the increased number of independent SNVs tested in sequence data.

We performed rare-variant burden tests using firth logistic regression in R, adjusting for the first 10 PCs, sex and platform(Firth, 1993; Heinze and Ploner, 2002). Variants for the rare-variant burden tests have been aggregated on multiple levels; gene, protein superfamilies, pathways, druggable categories and exome-wide. Genic regions were defined as all transcripts in the GRCh37.p13 version of Ensembl Biomart(Smedley et al., 2015). Higher level aggregation for burden analysis was performed by creating genesets. These genesets are based on: (a) protein superfamilies(Wilson et al., 2007); (b) drugable categories as defined by the drug-gene interaction database (Cotto et al., 2017); and (c) pathways downloaded from GSEA, using curated genesets v6.1 from KEGG, BioCarta or Reactome(Mootha et al., 2003; Subramanian et al., 2005).

We tested genes or genesets when we could identify ≥ 5 individuals with ≥ 1 variant. We used three definitions for ‘rare’: minor allele frequency cutoffs 1% and 0.5%. and variants not observed in ExAC(Lek et al., 2016). We classified variants based on their functional annotation (disruptive, damaging, missense-non-damaging, and synonymous, and described previously(Genovese et al., 2016)). Briefly, frame-shift, splice site, exon loss, stop gained, stoploss, startloss and transcription ablation variants were regarded as disruptive variants. We defined damaging variants as missense variants (resulting in an amino-acid change) predicted as damaging by *all* of seven methods: SIFT, Polyphen-2, LRT, Mutation Taster, Mutations Assessor, and PROVEAN(Genovese et al., 2016). Missense-non-damaging variants are missense variants that are not classified as damaging. Synonymous variants do not result in an amino-acid change. From these annotations, we created three variant sets for burden testing: (1) disruptive variants, (2) disruptive + damaging variants, (3) disruptive + damaging + missense-non-damaging variants. The synonymous category functions as a null category to check for biases when testing for association. We set the threshold for exome-wide significance in genic rare-variant burden analyses at p < 1.7 × 10^−6^. We acknowledge that this threshold does not fully account for the multiple testing burden introduced by the different variant sets, allele-frequency cut-offs, and various burden testing approaches.

### Data integration and annotation

After quality control, we performed functional annotation of all variants using snpEff V4.3T and SnpSift using the GRCh37.75 database (including Nextprot and Motif), dbSNFP v2.9, dbSNP b150 GRCh37pl3 and ClinVar GRCh37 v2.0(Cingolani et al., 2014; Gaudet et al., 2017; Landrum et al., 2016; Liu et al., 2016; Ruden, 2012; Sherry et al., 2001). We obtained population frequency estimates from gnomAD(Lek et al., 2016). To visualise the variant-level coverage from Project MinE and external sources, we included coverage information from Project MinE samples, gnomAD database (123,136 exome sequences plus 15,496 genome sequences). We further integrated tissue-specific gene expression profiles for 53 tissues from the GTEx resource (https://gtexportal.org/home/datasets) (GTEx Consortium, 2015). Finally, the available literature on each gene is presented through an iframe linking to either PubMed, UCSC, GeneCards, Ensembl, WikiGenes, GTEx or the GWAScatalog.

#### Existing ALS datasets

The browser also includes freely-available summary-level data for the 2016 ALS GWAS for download. Additionally, downloadable SKAT and SKAT-O burden testing results from 610 ALS cases and 460 controls with Chinese ancestry (Gratten et al, 2017) are available.

#### Language

The data browser can be accessed at http://databrowser.projectmine.com/. The interface is based on the statistical programming language R (v3.4.1, https://www.r-project.org/) together with the interactive web application framework Shiny (v1.0.5, https://shiny.rstudio.com/). Interactive visualisations have been created using base R and the Plotly library (v4.7.1.https://plot.ly/r/). The code is open-source and can be downloaded from https://bitbucket.org/ProjectMinE/databrowser.

### Informed consent

All participants gave written informed consent and the relevant institutional review boards approved this study. The informed consent clearly indicates that there is no duty to hunt for clinically actionable results and that participants will not be re-contacted for genotyping results.

## Results

### Dataset

The databrowser currently comprises 4,366 ALS cases and 1,832 age- and sex-matched controls whole-genome sequenced and quality control processed as part of the broader Project MinE effort.

### Quality control and association analysis

The quality controlled dataset includes 6,198 individuals and describes more than 105 million SNVs and indels. In this sample we have limited power to detect genome-wide significant association in a single variant framework and as a result we did not find any variants reaching genome-wide significance. In our rare-variant burden framework we find that the excess of disruptive and damaging variants at MAF < 1% in the canonical transcript of *NEK1* in ALS patients compared to controls reaches exome-wide significance (p = 2.31 × 10^−7^, odds ratio = 3.55 [95% confidence interval = 2.02 − 6.26], **Fig. 2**). We also noticed that some genes might contain a transcript specific burden, most notably in TARDBP (**Supplementary Table 3 and Supplementary Fig. 12**).

**Figure 2.**
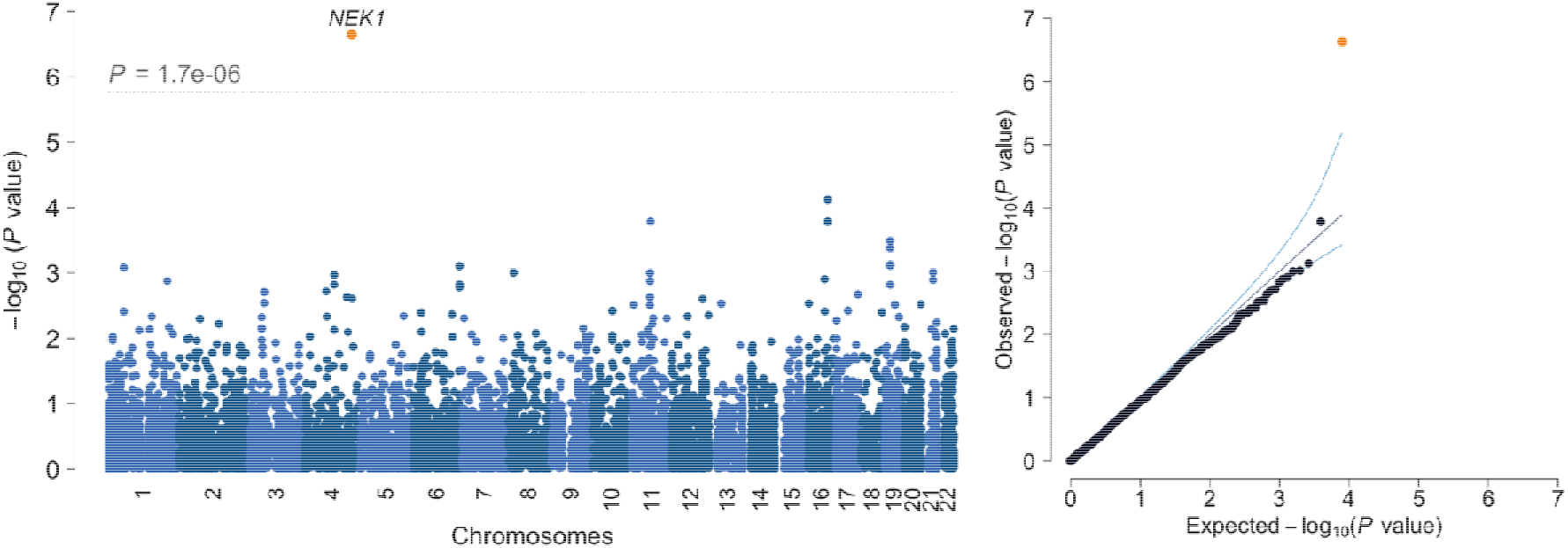
Manhattan and QQ-plot. Results are shown for genic (canonical transcripts only) firth logistic regression including variants with a MAF<1% and categorised as disruptive and damaging, *λ*_GC_ = 0.907, *λ*1000 = 0.964.

Next, we aggregated all variants across the exome. We observed no difference in the exome-wide burden of *synonymous* variants between cases and controls, which provides no indication for systemic confounding of burden analyses using higher-order variant aggregation strategies. Therefore, we proceeded to test a genome-wide excess of rare *non-synonymous* variants among ALS patients. In contrast to similar analyses in schizophrenia (Genovese et al., 2016) and educational attainment (Ganna et al., 2016), we found no evidence for such excess in any variant set combining all allele frequency cut-offs and functional classification.

Furthermore, we do not find any protein families, druggable categories significantly enriched for rare variants after collapsing allele-frequency cut-offs and variant classification. All association analysis results are available for download at the browser website.

### Databrowser

By entering a gene or transcript in the databrowser you will be shown a visualisation of the rare-variant burden tests, as well as several other components (**Fig. 3**).

**Figure 3.**
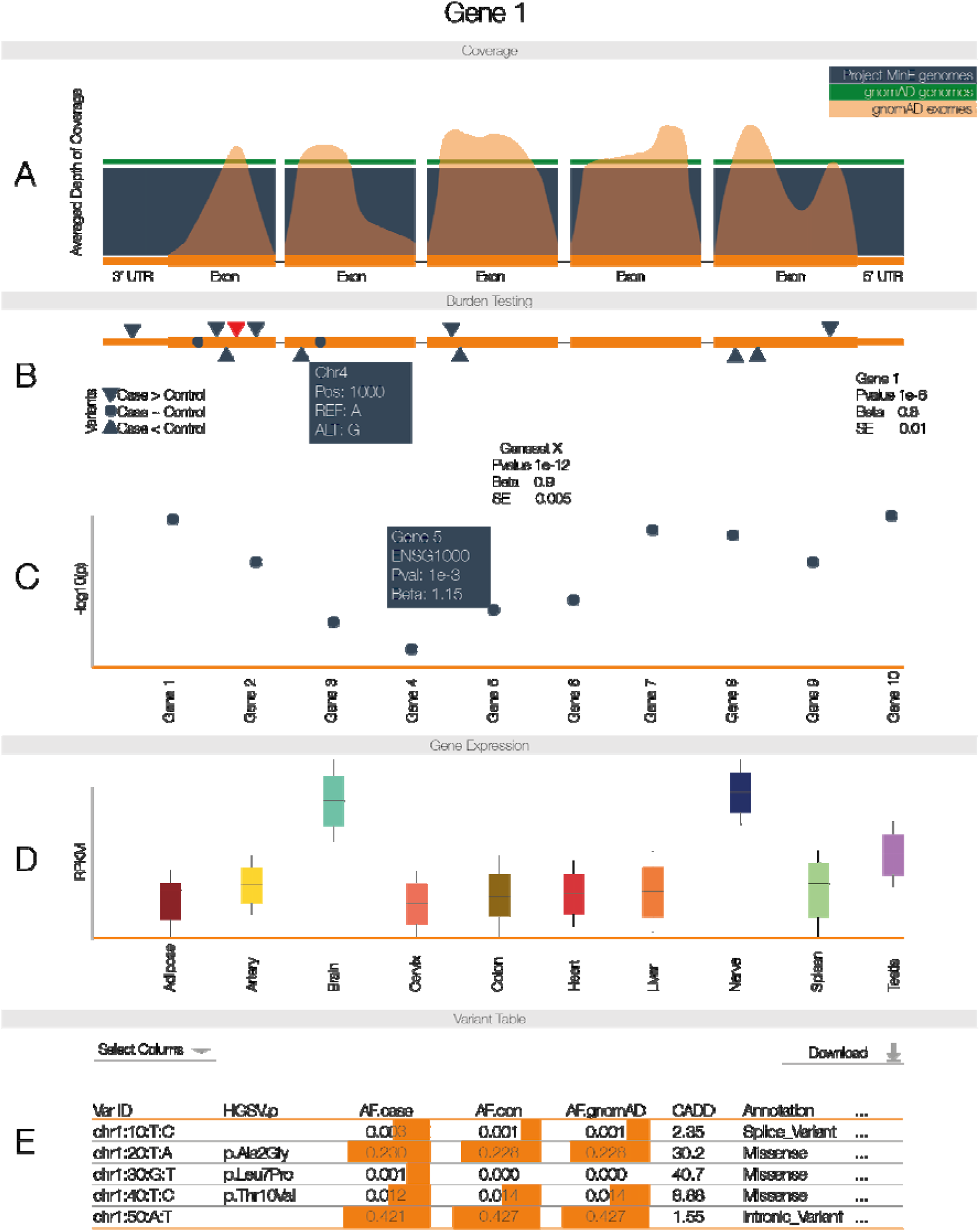
Databrowser. After entering the gene name (HGNC, Ensembl gene (ENSG) or transcript (ENST) identifier) in the search box on the homepage, you will be directed to the gene-specific page. **A** Averaged depth of coverage in the Project MinE dataset, compared to public data and indicating quality of coverage in the region. B Firth logistic regression-based genic burden tests. Triangles indicate variant locations. Red triangles reach nominal significance in the single variants association test. Hovering over the triangles to obtain more information about that variant. **C** Firth logistic regression-based geneset burden test. Tests are based on pathways, gene families or druggable gene categories. To elucidate the gene or genes genes driving a signal in the geneset, a Manhattan plot indicates the genic burden results for each of the genes included in the geneset. Hovering over individual genes will reveal more information about that gene. **D** Gene expression profiles extracted from GTEx. **E** Variant table. By default, a subset of variant information is shown; columns of interest can be selected from the dropdown menu. Minor allele frequency is based on all unrelated and QC passing samples in the Project MinE dataset (6,198 genomes). Frequency information is also stratified by phenotypic status and compared to public exome and whole genome data. For comparison, we have indicated the allele frequency on a log scale with orange bars; the longer the bar, the higher the allele frequency. Variant filtering can be customised using the search boxes below the header of each column. All data, including case/control frequencies, are available for download in a tab-delimited file. For a more detailed view of the databrowser, see **Supplementary Fig. 11**.

#### Transcript details

Here we describe the elementary transcript details for the gene of interest. This includes the Ensembl transcript ID, Ensembl Gene ID, number of exons and genomic coordinates as described in the GRCh37 build.

#### Coverage information (Fig. 3a)

To illustrate whether a particular gene/transcript or exon has been adequately covered to detect variation, we have included a graphical representation of average depth of coverage. This graph also includes the coverage information from the ExAC database to illustrate the difference in coverage between genome- and exome-sequencing. Optionally, the coverage across introns can be visualised.

#### Genic burden results (Fig. 3b)

Burden testing, by definition, aggregates many variants. This approach can increase statistical power to find an association, but can obscure which variant(s) are driving a potential association. Therefore, we have included an interactive graphical representation of the gene indicating where variants are located and whether these variants are case or control-specific. Hovering over a specific variant will reveal the position, alleles, heterozygous and homozygous allele counts in ALS cases and controls, and functional annotation of the variant. We additionally provide the burden test statistics. To further facilitate interpretation, we describe the burden test properties and relevant references in a dropdown menu “Burdentest Info.” We have performed genic burden results for all transcripts.

#### Geneset burden results (Fig. 3c)

Here, we show burden test results for genesets such as protein families and druggable targets to which the selected gene belongs. This includes a mini-Manhattan plot generated to indicate which genes might be driving an association signal in the geneset by plotting their individual genic burden results.

#### Tissue-specific gene expression (Fig. 3d)

This panel shows gene expression levels across all general tissues included in GTEx.

#### Variant annotation table (Fig. 3e)

Each variant has been extensively annotated and aggregated in a customizable table. By default, only allele frequency in cases and controls, comparison to gnomAD genomes and exomes, and amino acid change, impact and functional consequence are shown. All information can be downloaded in tabular form.

#### Gene-specific literature

To provide background information on the gene’s function and disease association from literature, we have included an iframe linking to PubMed, UCSC, GeneCards, Ensembl, WikiGenes, GTEx and the GWAScatalog. This allows a user to rapidly extract information from various resources while staying on the same page.

### Group and individual level data sharing

The summary statistics for the latest GWAS, WGS single variant association and all WGS burden analyses can be downloaded directly. Access to individual-level data can be requested by providing a digital form with a brief research proposal (https://www.projectmine.com/research/data-sharing/).

### Duplicate and relatedness checks

We have created sumchecks for each individual in our dataset. Sumchecks are hashes which have been created, based on a small subset of SNPs, which allow for the identification of duplicates without sharing the genetic data itself. If researchers wish to check duplicates with our dataset, they can simply request the sumchecks, create hashes for their own data and compare the hashes. The code to generate the hashes and the list of SNPs used is available on the Project MinE Bitbucket. These hashes only identify duplicate samples, and in some instances relatedness information can be valuable, e.g., extending pedigrees or meta-analyses. Therefore, we will perform the relatedness checks when a statement is uploaded that this information will be used for academic purposes only and will not be used to re-identify individuals without consent. These checks do not require a data-access request nor approval.

### Technical details

The whole website, including data storage, runs on a dual core server with 4Gb RAM and needs <50Gb of storage. As of July 2018, we have had over 6,200 sessions from over 1,500 users.

## Discussion

Both research and clinical work increasingly rely on open-access databases to find newly-associated variants and interpret genetic findings when counselling patients(Bonàs-Guarch et al., 2018). Therefore, sharing de-identified data is instrumental to ensuring scientific and clinical progress, and patient-derived data should not be regarded as intellectual property nor as trade secret (Directors, 2017) (“Data sharing and the future of science,” 2018). Also, most genetic browsers are based on healthy individuals, or unselected individuals who might carry specific rare genetic variants which hampers adequate comparison to a sample of patients from another geographical region. With exactly this in mind, we developed a unique, publicly-available, disease-specific databrowser which serves as a transparent framework for sharing data and results in ALS genetics. The Project MinE Databrowser contains an unprecedented amount of WGS data from ALS patients, more than doubling the currently-available exome based databases, and provides (meta)data in far greater detail. The intuitive design facilitates interpretation of robust statistical association analyses by presenting detailed metadata and through integration with population-based observations, biological/functional context and literature. As a result, we make our data and results accessible to a broad public of diverse backgrounds and for any research initiative. The databrowser provides an easy framework for other consortia who are generating similar genetic data and results in ALS and other diseases.

The data has already provided a backbone for new gene discovery and variant interpretation in ALS. For example, subsets of the current dataset have been incorporated in previous publications which identified *C21orf2, NEK1* and *KIF5A* (Kenna et al., 2016; Nicolas et al., 2018; van Rheenen et al., 2016). The resource will continue to grow as the Project MinE consortium does, and will thus increasingly allow for more reliable identification of true positives (Project MinE ALS Sequencing Consortium, 2018; Van der Spek et al., 2018). The growth in both sample size and ancestral diversity will increasingly reflect the ALS mutation spectrum and yield increasingly accurate estimations of effect sizes in the general population. The browser can also offer researchers quick, easy to access to a reliable dataset for significant improvement in statistical power without financial burden.

One of the major goals of the databrowser is to allow cross-disciplinary interrogation and interpretation of the data with minimal effort. We enable this through the intuitive display of individual variant level data, statistical results and through the integration with databases including GTEx and gnomAD. The databrowser ensures transparency and continued reevaluation of established associations, vitally important for clinical laboratories to make appropriate variant classifications(Project MinE ALS Sequencing Consortium, 2018). Furthermore, we aim to facilitate the design of functional experiments by showing which variants, might be driving a genic burden signal and if these are located in specific exons and therefore specific protein domains.

Project MinE is largely crowd-funded and the ALS-community is highly engaged in the scientific progress in our field. Consequently, we feel an obligation to give something back to the community and promote data sharing in general. We hope that our databrowser will inspire similar efforts in other fields. The Project MinE databrowser is a light-weight and open-source R script that can easily be adapted to serve other consortia and thus share similarly important data. Further, we aim to improve data sharing by encouraging fellow researchers to gain access to individual-level data by submitting an analysis proposal to the consortium. After access is granted, analyses can be performed on the compute facilities of SURFsara, a supercomputer based in Amsterdam, The Netherlands. Researchers will only need to pay a minimal fee to compensate costs for their core hours and data storage requirements.

Project MinE continues to work forward to its ultimate goal of whole-genome sequencing 15,000 cases and 7,500 matched controls, as well as combining the data with publicly-available control data. Current efforts also focus on single SNV and aggregated SNV analyses of autosomal chromosomes. Future efforts will aim to include sex chromosomes, indels, structural variation (in particular, repeat expansions(Dolzhenko et al., 2017)) and noncoding burden analyses. Additionally, Project MinE is collecting methylation data on all samples using the Infinium Human Methylation 450K and EPIC BeadChip. These data and analyses will also be shared expeditiously through our databrowser prior to publication. As the project proceeds and data generation continues apace, we intend for the browser to pave the way for more accurate diagnosis and prognosis, aid in the identification of novel disease-associated genes, and elucidate potential novel therapeutic targets.

## Acknowledgements

This work was carried out on the Dutch national e-infrastructure with the support of SURF Cooperative.

## Code availability

Source-code for the databrowser is available at https://bitbucket.org/ProjectMinE/databrowser

## Author contributions

R.A.A.v.d.S wrote source-code for the databrowser and together with W.v.R. and J.H.V. designed the databrowser, performed and discussed all analyses and wrote the manuscript. S.P performed QC on the WGS dataset and was involved is revising the manuscript. K.P.K, R.L.McL, M.M., A.D., G.T. contributed to data collection and discussions to improve the design of the databrowser. L.H.v.d.B. advised and assisted in study design. Unmentioned authors contributed to data collection and funding in their countries.

