## Supplementary Information for "The Project MinE databrowser: bringing large-scale whole-genome sequencing in ALS to researchers and the public"

### Supplementary Materials and Methods

Data merging and initial site filtering

Per individual, WGS data was stored in both BAM and Illumina gVCF format. These gVCFs contain single-nucleotide variants (SNVs), short insertions and deletions (indels), and large structural variants (SVs). To begin quality control, we merged all sample into a single file using the Illumina ‘agg’ tool v0.3.4 (https://github.com/Illumina/agg). ‘Agg’ first generates metadata across all samples, typically batched into groups (e.g., n = 50) to minimise CPU time, and then proceeds to extract genotypes for all samples across all sites containing at least one non-reference allele in the full case-control dataset. This process results in a VCF of all samples and all variants with minor allele count ≥ 1.

The resulting merged VCF contained all possible sites, regardless of whether or not they passed the Isaac pipeline set of variant filters. We therefore applied basic site filtering to the initial merged VCF. Specifically, we set sites with a genotype quality (GQ) < 10 to missing using bcftools and single-nucleotide variants (SNVs) and indels with quality (QUAL) scores < 20 and < 30, respectively, were removed. We then removed variants with missingness > 10% (typically induced by setting genotypes with GQ < 10 to missing). To ensure unique marker identifiers at all sites, particularly for those multiallelic sites that were devolved by processing the data through ‘agg,’ we labeled all variants using the following nomenclature: chromosome:position:reference_allele:alternate_allele.

#### Sample-level quality control

We fixed genotype ploidy on the sex by first inferring biological sex from the available SNP array data, and then using the ‘fix-ploidy’ option in bcftools.

We then performed sample-level quality control (QC). We calculated the transition-transversion ratio in each sample using SnpSift 4.3p (**Supplementary Fig. 1**). In WGS data, the expected transition-transversion ratio is ~2.0; a number much lower than this (i.e. approaching 0.5, in accordance with the expected number of transitions and transversions if genotypes were called randomly, **Supplementary Fig. 1**) indicates an enrichment for false-positive genotype calls. We removed two samples with a Ti/Tv ratio > 6 standard deviations (sd) from the full distribution of samples.

Per sample, we also calculated (a) the total number of SNVs, (b) total number of indels, and (c) total number of singletons (**Supplementary Fig. 1**). We removed samples with a total number of SNPs > 6 sd from the mean. The shift in sequencing platforms from HiSeq 2000 to HiSeq X (which occurred in parallel with a change in the calling pipeline, to improve indel detection) caused an obvious shift in observed indels per sample. Samples were thus filtered by platform (HiSeq 2000 or HiSeq X) and removed samples with number of indels > 6 sd from the mean of their respective group. Finally, we identified samples with an excess number of singletons (calculated by cohort, to avoid overly-stringent filtering due to population stratification); samples with a total number of singletons > 6 sd from the sample distribution were removed.

Next, we calculated sample-level missingness and removed samples with > 5% missingness (**Supplementary Fig. 2**). We calculated average sample depth and again observed noticeable differences between those samples sequenced on the HiSeq 2000 and the HiSeq X, where average depth of coverage was somewhat higher (35X, on average) for samples sequenced on HiSeq 2000 compared to the samples sequenced on the HiSeq X (25X, on average). We removed no samples at this step. We then subsetted the sequence data down to those markers that overlapped with the 2.5M array genotyping data. Across the intersect of markers, we calculated the sample concordance between the sequence and array data, and removed all samples with concordance < 96% (**Supplementary Fig. 2**).

Using X chromosome variants, we tested to see if biological sex (inferred from the X chromosome data) was concordant with the sex as annotated in the available phenotype information (**Supplementary Fig. 3**). We excluded 62 (of 6,579) samples with mismatching information.

We performed the remaining sample QC on a high-quality set of ~100,000 autosomal variants with: minor allele frequency (MAF) > 10%; genotype missingness < 0.1%; residing outside four complex regions (the major histocompatibility complex (MHC) on chromosome 6; the lactase locus (*LCT*), on chromosome 2; and inversions on chromosomes 8 and 17); excluding A/T and C/G variants. We used this set of markers to calculate inbreeding in two ways: first, by calculating inbreeding coefficients using Plink 1.9 (--het, **Supplementary Fig. 3**); and secondly, by calculating the ratio of heterozygous to homozygous non-reference genotypes per sample. In the first instance, we removed samples > 6 sd from the full sample distribution (**Supplementary Fig. 3**). In the second instance, we filtered samples for inbreeding coefficients on a cohort-by-cohort basis and excluded individuals > 6 sd from the cohort distribution (**Supplementary Fig. 4**).

We estimated kinship coefficients (i.e., relatedness) using the KING method, as implemented in the SNPRelate package in R. As samples were ascertained from a number of countries, we used the KING method, as it calculates kinship in the presence of potential population stratification (a potential confounder in other identity-by-descent approaches, such as that implemented in Plink). In some instances, research groups had intentionally ascertained related samples. We identified all pairs of related individuals (kinship > 0.0625). Of these, several pairs included one sample appearing to be related to several other samples in the data (likely due to sample contamination; **Supplementary Fig. 3**). Samples related to > 100 other samples in the data were dropped; true related pairs were left in the data. For burden testing, we excluded these related samples (kinship coefficient > 0.0625). For single variant association analysis, we used a linear mixed model in GCTA, including a genetic relationship matrix and the first 20 principal components (PCs), alleviating the need to exclude related samples.

Lastly, we used principal component analysis (PCA) implemented in EIGENSTRAT to visualise potential structure in the data, induced by population stratification or other variables (**Supplementary Fig. 4**). Projection onto the HapMap 3 populations indicated that the samples were primarily of European ancestry, though some were of African or East Asian ancestry, while other samples appeared to be admixed. PCA across the dataset alone revealed structure induced not only by population but also by sequencing platform/calling algorithm (e.g., principal component 2, **Supplementary Fig. 5H**). However, because of a balanced case-control ratio in both batches, we observed a very small effect of platform/calling algorithm when including it as a covariate in association testing. A summary of all sample QC, including thresholds and removed samples, is provided in **Supplementary Table 2**.

Variant-level quality control

To clean variants, we first inferred a set of QC thresholds from the set of SNVs falling on chromosomes 1-22 and then extrapolated these thresholds to filter all variants, including indels and variants on the sex and mitochondrial chromosomes.

We calculated Hardy-Weinberg equilibrium (HWE) in controls only, on a cohort-specific basis (to avoid potential population confounding) and removed all variants with HWE p < 1 × 10^-6^. We calculated differential missingness between cases and controls and removed any variants with p < 1 × 10^-6^.

Next, we binned the variants by a number of metrics: depth of coverage, minor allele frequency, missingness, quality (QUAL) score, and passing rate (**Supplementary Fig 7-8**). The last metric, passing rate, indicates the proportion of samples for which the variant was annotated as ‘PASS’ing variant filters in the original, per-sample gVCF data. For example, a passing rate of 70% indicates that a variant is annotated as ‘passing’ the Isaac thresholds in 70% of all samples.

Once we had stratified the variants by these metrics, we calculated (for each bin) the transition/transversion (Ti/Tv) ratio and the ratio of heterozygous to homozygous non-reference genotypes (het/hom-non-ref) and then plotted the bins according to these metrics (**Supplementary Fig. 7-8**). From these visualizations of the data, we could infer the following QC thresholds and remove: variants with total depth < 10,000 reads (i.e., ~1.53X per sample) or > 226,000 reads (i.e., ~34.8X per sample), variants with missingness > 5%, and variants with a passing rate < 70%. We did not filter variants on minor allele frequency or QUAL score.

Scripts used for performing data merging and sample- and variant-level quality control are available through the Project MinE BitBucket (<https://bitbucket.org/ProjectMinE/databrowser>). Scripts include calls to PLINK, bcftools, SnpSift, EIGENSTRAT, and SNPRelate, as well as the relevant command-line options used for the QC steps described here.

### Supplementary tables and figures

| **Country** | **N** |
| --- | --- |
| Belgium | 568 |
| Ireland | 408 |
| Netherlands | 2,995 |
| Spain | 359 |
| Turkey | 224 |
| United Kingdom | 1,468 |
| United States | 557 |
| Total | 6,579 |

Supplementary Table 1. Total number of samples before quality control, stratified by country.

|  | Belgium | Ireland | Netherlands | Spain | Turkey | UK | USA | Total |
| --- | --- | --- | --- | --- | --- | --- | --- | --- |
| Total SNPs | 0 | 1 | 17 | 2 | 0 | 6 | 11 | 37 |
| Total indels | 1 | 1 | 12 | 2 | 0 | 6 | 10 | 32 |
| Total singletons | 1 | 1 | 2 | 1 | 0 | 1 | 1 | 35 |
| Ti/Tv ratio | 1 | 0 | 1 | 0 | 0 | 0 | 0 | 2 |
| Missingness | 2 | 0 | 0 | 0 | 0 | 0 | 0 | 2 |
| Geno-Seq concordance | 0 | 0 | 4 | 0 | 0 | 0 | 0 | 4 |
| Depth of cov | 0 | 0 | 0 | 0 | 0 | 0 | 0 | 0 |
| Relatedness | 0 | 1 | 23 | 1 | 1 | 15 | 2 | 43 |
| Het/hom-non-ref ratio | 0 | 1 | 15 | 1 | 1 | 7 | 1 | 26 |
| Inbreeding | 3 | 1 | 17 | 1 | 1 | 7 | 1 | 31 |
| Sex check | 13 | 1 | 17 | 15 | 8 | 7 | 1 | 62 |
| PCA | 0 | 0 | 0 | 0 | 0 | 0 | 0 | 0 |
| Total failures including sex check | 18 | 3 | 49 | 16 | 9 | 25 | 13 | 133 |
| Total failures excluding sex check | 6 | 2 | 41 | 2 | 1 | 20 | 12 | 84 |

Supplementary Table 2. Quality control. Summary of the number of samples that have failed quality control steps. Please note that some samples fail multiple steps; therefore, the sum of a column will not be equal to the number of samples that actually failed.

| **Ensembl**  **Transcript ID** | **variants** | **Controls*** | **Cases*** | **P** | **Beta** | **SE** |
| --- | --- | --- | --- | --- | --- | --- |
| ENST00000439080 | 19 | 0 | 28 | 3.4 × 10^-5^ | 3.53 | 1.64 |
| ENST00000240185^C^ | 21 | 6 | 37 | 0.017 | 0.93 | 0.44 |
| ENST00000315091 | 5 | 6 | 12 | 0.71 | -0.19 | 0.50 |
| ENST00000473118 | 5 | 6 | 12 | 0.71 | -0.19 | 0.50 |
| ENST00000473869 | 5 | 6 | 12 | 0.71 | -0.19 | 0.50 |
| ENST00000476201 | 5 | 6 | 12 | 0.71 | -0.19 | 0.50 |
| ENST00000472476 | 3 | 0 | 3 | - | - | - |
| ENST00000477447 | 0 | - | - | - | - | - |
| ENST00000496840 | 0 | - | - | - | - | - |

Supplementary Table 3. Transcript-specific burden for TARDBP. All non-synonymous variants with a MAF < 0.5% were included. Firth logistic regression was performed with the first 10PCs, sex and platform were included as covariates. Transcript ENST00000439080 clearly shows an increased burden as compared to the other transcripts. **Supplementary Fig. 12** shows the burdenplot of the first 3 transcripts. ^C^ Canonical transcript. * Number of controls or cases carrying at least 1 variant.

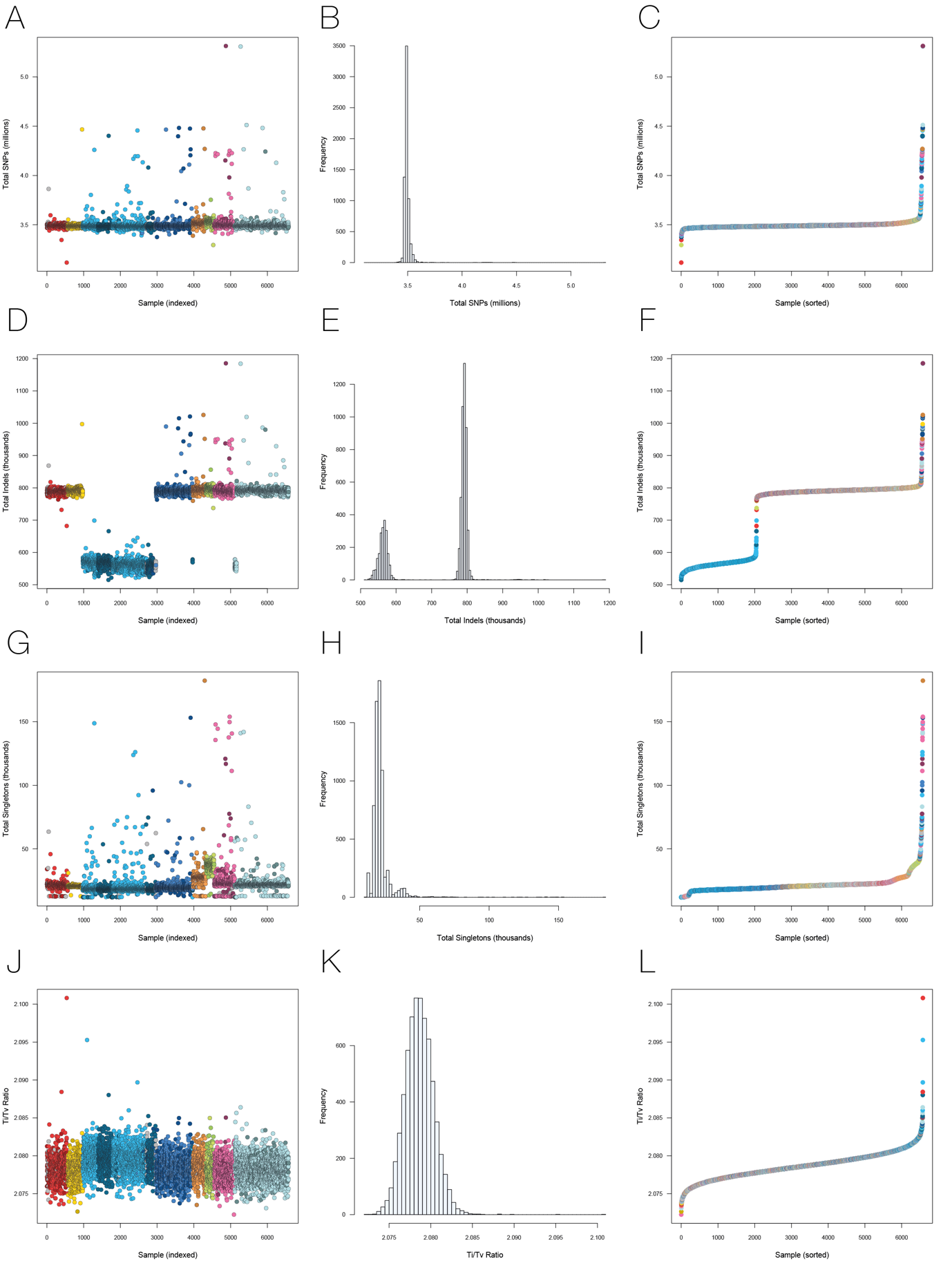

Supplementary Figure 1. Individual-level quality control part 1. Panel **A**,**B**,**C** Total number of SNPs per samples. Mean = 3.5M, SD = 70K, filtering threshold = 6SD. Panel **D**,**E**,**F** Total number of Indels. Mean_HiSeq200_ = 560K, SD_HiSeq2000_ = 13.5K, filtering threshold_HiSeq2000_ = 6SD. Mean_HiSeqX_ = 792K, SD_HiSeqX_ = 17.4K, filtering threshold_HiSeqX_ = 6SD. Panel **G**, **H** and **I** Total number of Singletons. Outliers were determined per cohort. Please see **Supplementary Fig. 4**. Panel (**J**,**K**,**L**) Ti/Tv ratio. Mean=2.08, SD = 0.002, Filtering threshold = 6SD.

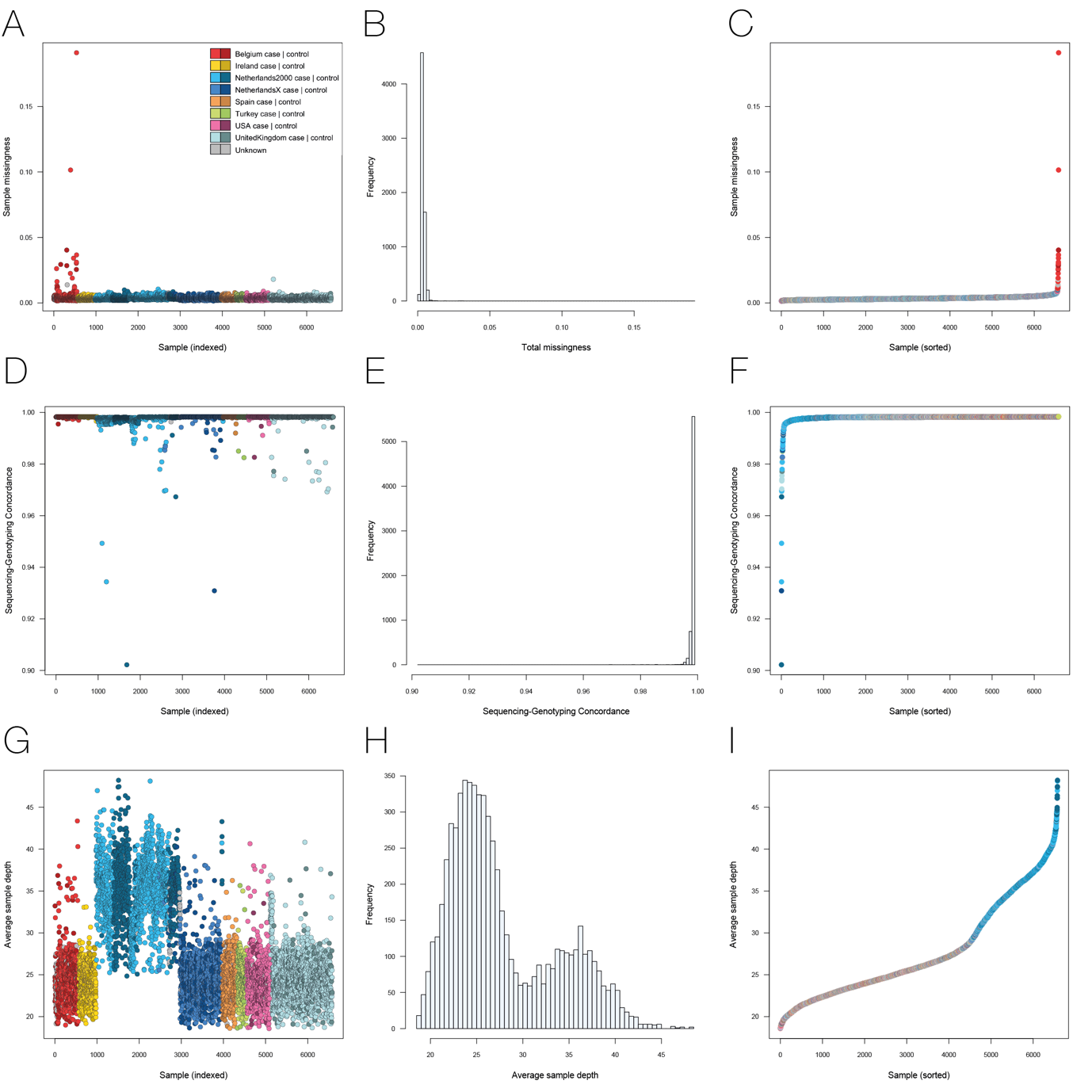

Supplementary Figure 2. Individual-level quality control part 2. Panel **A**,**B**,**C** Sample Missingness. Filtering threhold is set at 5%. Panel **D**,**E**,**F** Sequencing vs genotyping concordance (Illumina 2.5M). Filtering threshold is set at 96%. **G**,**H**,**I** Averaged sample depth. There is a clear difference in depth of coverage when comparing HiSeq2000 data with HiSeq X data. Please note that HiSeq2000 data consists of 100bp reads while HiSeq X consists of 150bp reads.

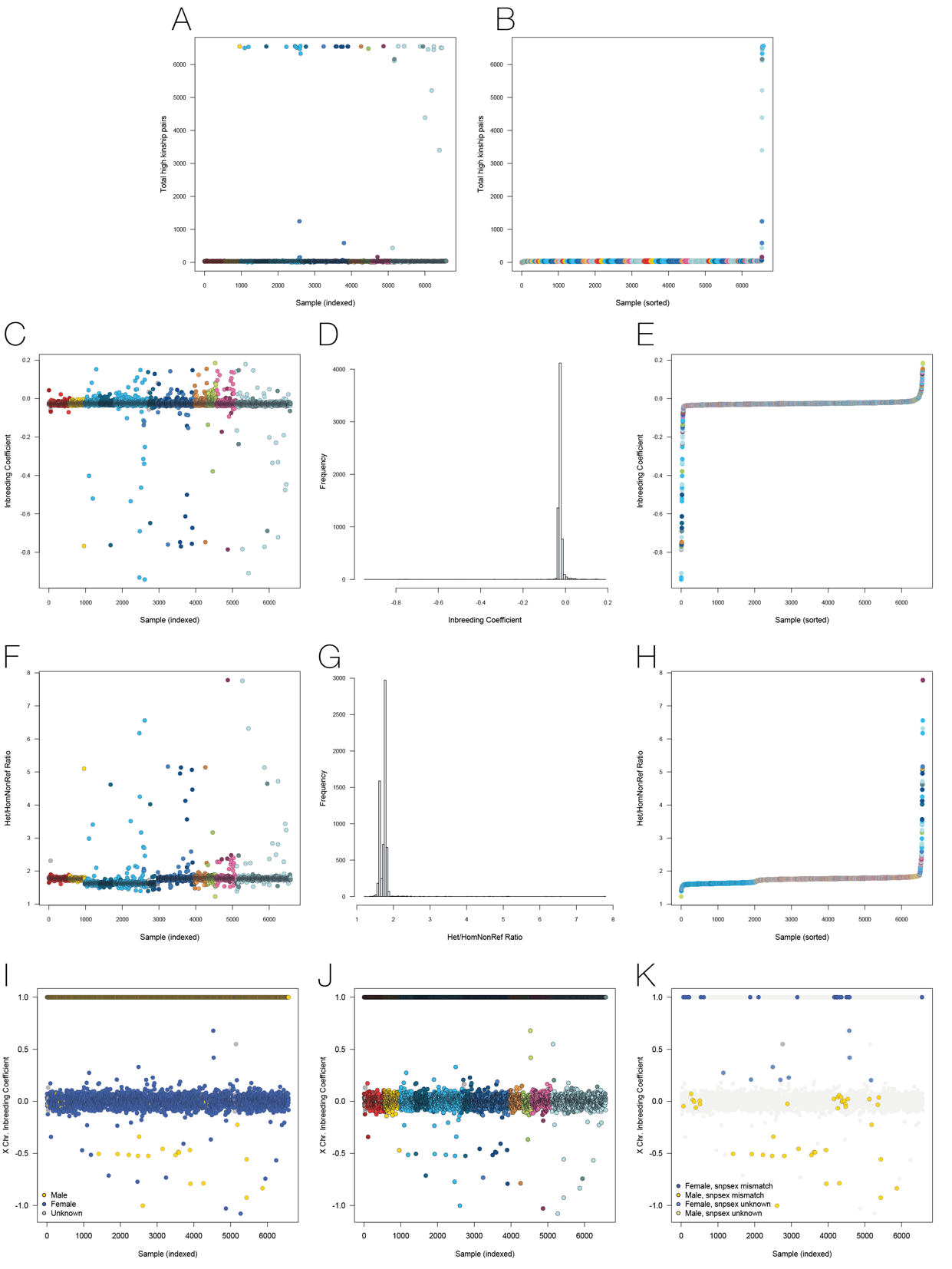

Supplementary Figure 3. Individual-level quality control part 3. **A**,**B** Relatedness as calculated using KING. Samples found in >100 relatedness pairs (kinship > 0.0625) were excluded from further analyses. Please note that this does not exclude all related samples. For single variant association analyses, related samples are retained and corrected for using a genetic relationship matrix (GRM). For burden analyses, additional samples are excluded based on relatedness (kinship > 0.0625). **C**,**D**,**E** Inbreeding coefficients for all samples. Samples are filtered by cohort. **F**,**G**,**H** Het/hom-non-ref ratio. Mean = 1.74, SD = 0.228, filtering threshold is 6SD. **I**,**J**,**K** Seks check based on chromosome X inbreeding coefficient. Excluded samples are indicated in K.

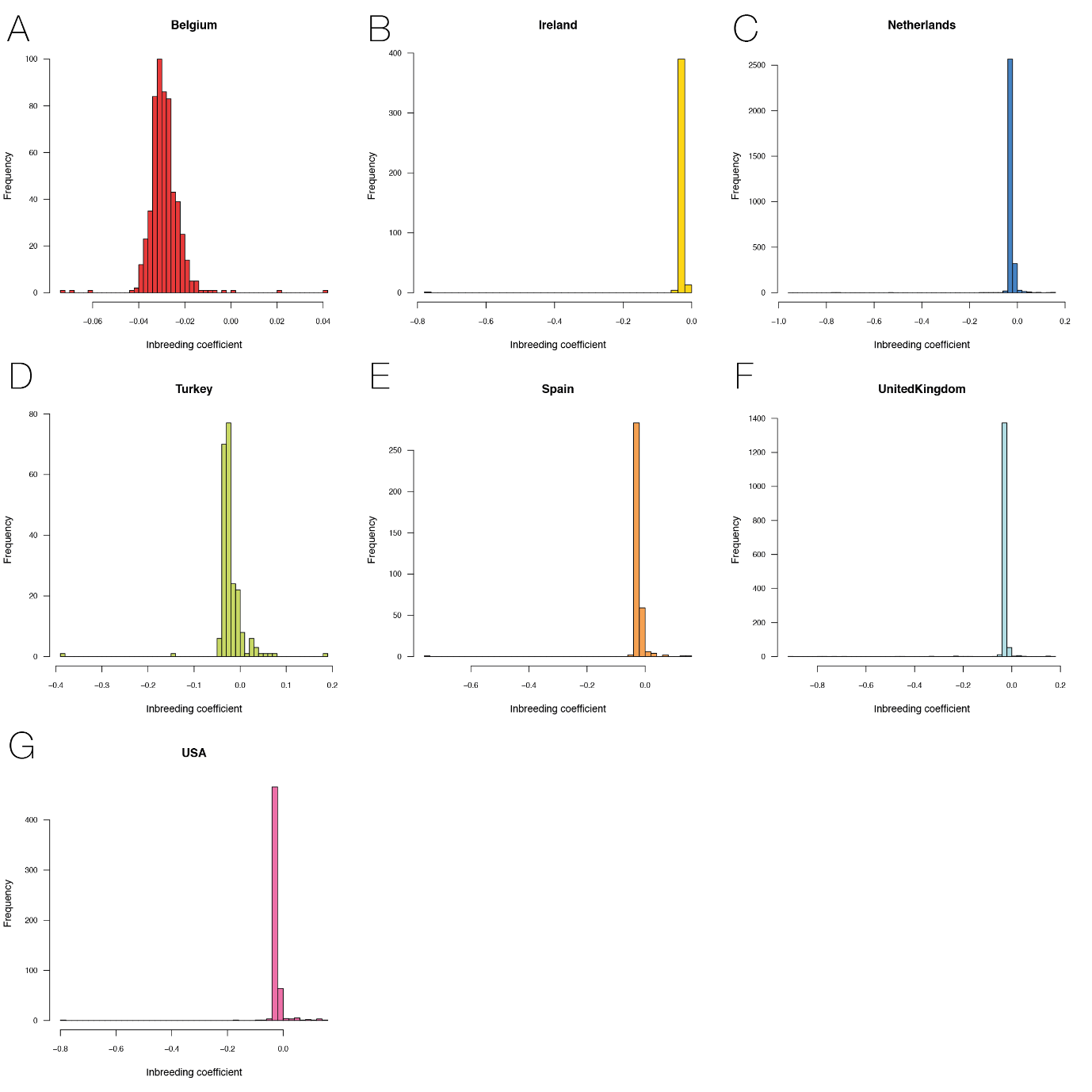

Supplementary Figure 4. Inbreeding coefficient per cohort. For each cohort samples with >6SD are excluded from further analyses.

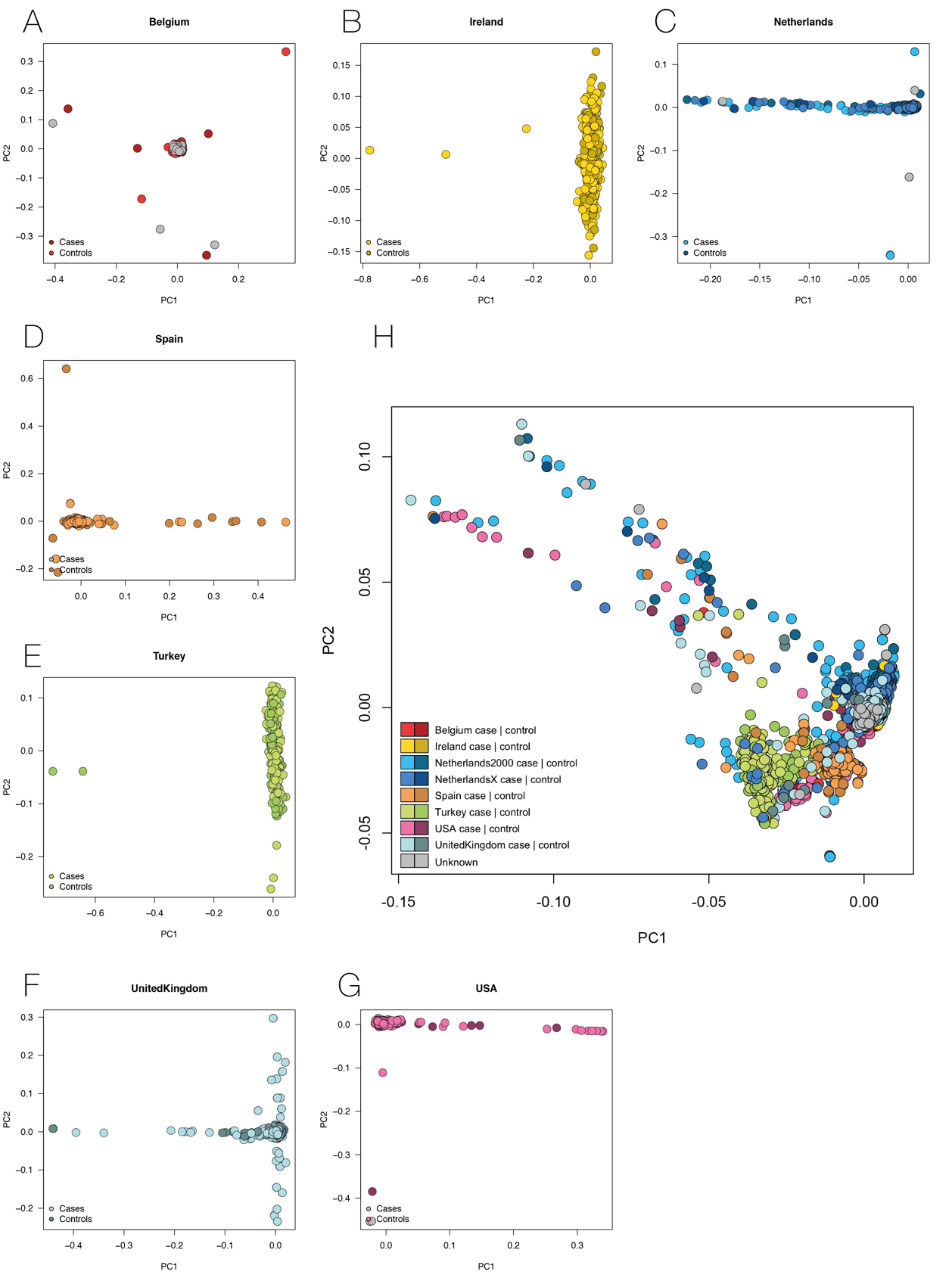

Supplementary Figure 5. Principal component analysis. **A-G** PC1 vs PC2 per country of origin. **H** PC1 vs PC2 over all cohorts. Please note the Netherlands has been split into two batches: the HiSeq2000 data and the HiSeq X data. In principle component space, these batches segregate from each other. However, because of a balanced case-control ratio in both batches, we observe a very small effect of platform when adding it as a covariate in association testing.

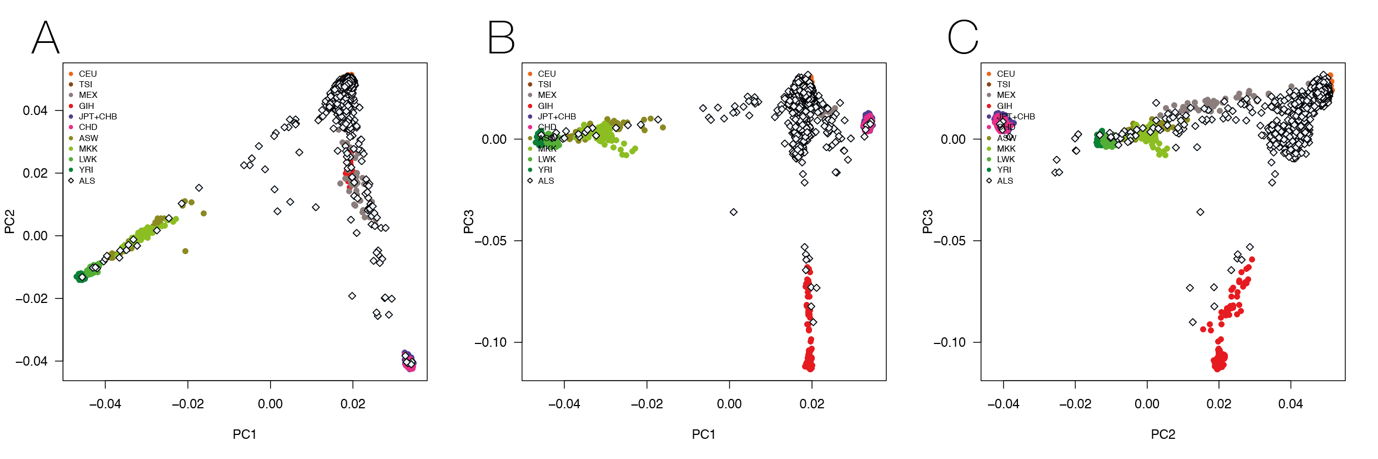

Supplementary Figure 6. PCA projected to HapMap

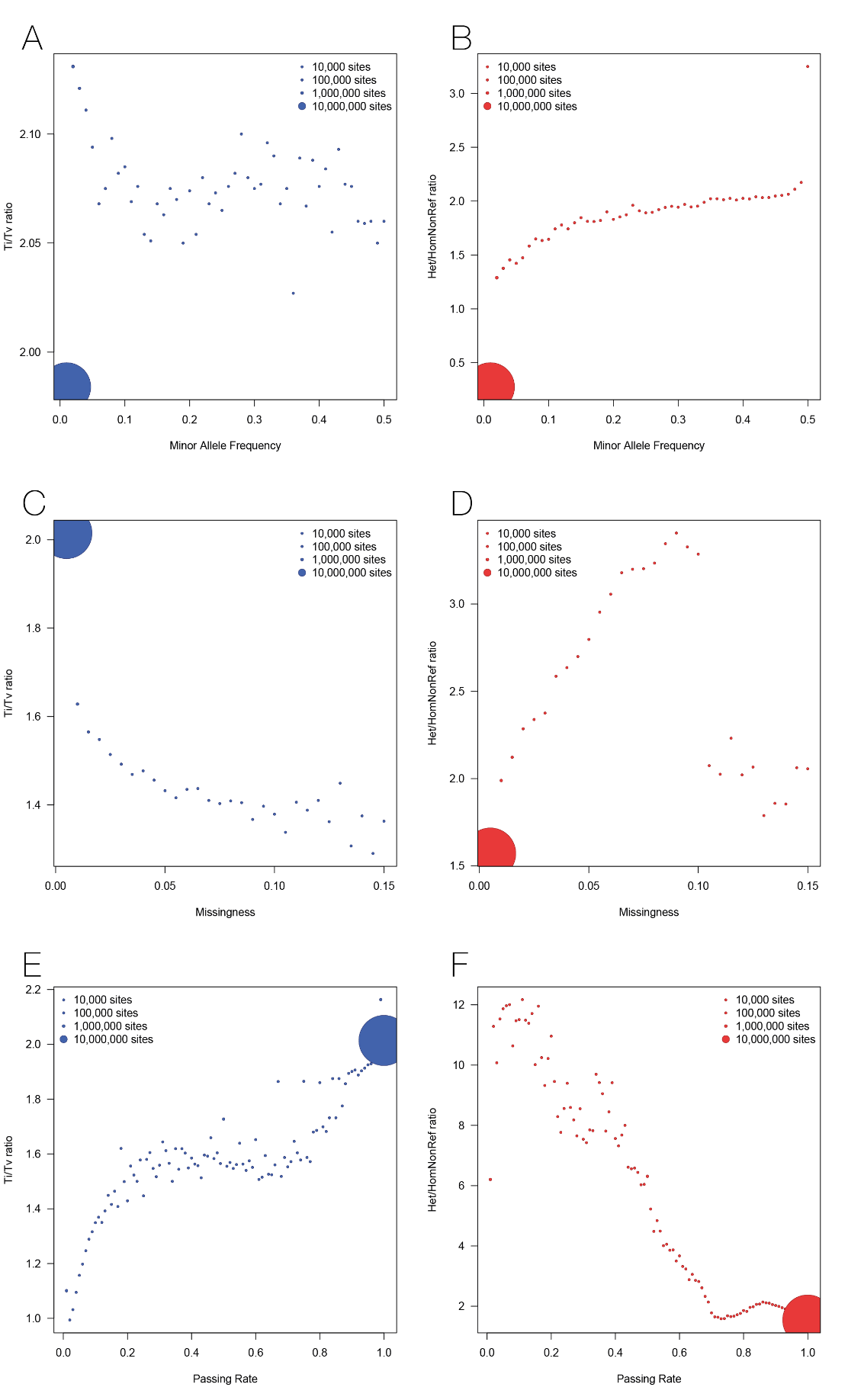

Supplementary Figure 7. SNP-level quality control part 1. Ti/Tv ratio and Het/Hom-non-reference ratios for MAF bins **A**,**B**, Missingness bins **C**,**D** and QC Passing Rate **E**,**F**. From these visualizations of the data, we could infer the following QC thresholds: variants with missingness > 5% were removed, and variants with a passing rate < 70% were also removed. Variants were not filtered based on either minor allele frequency.

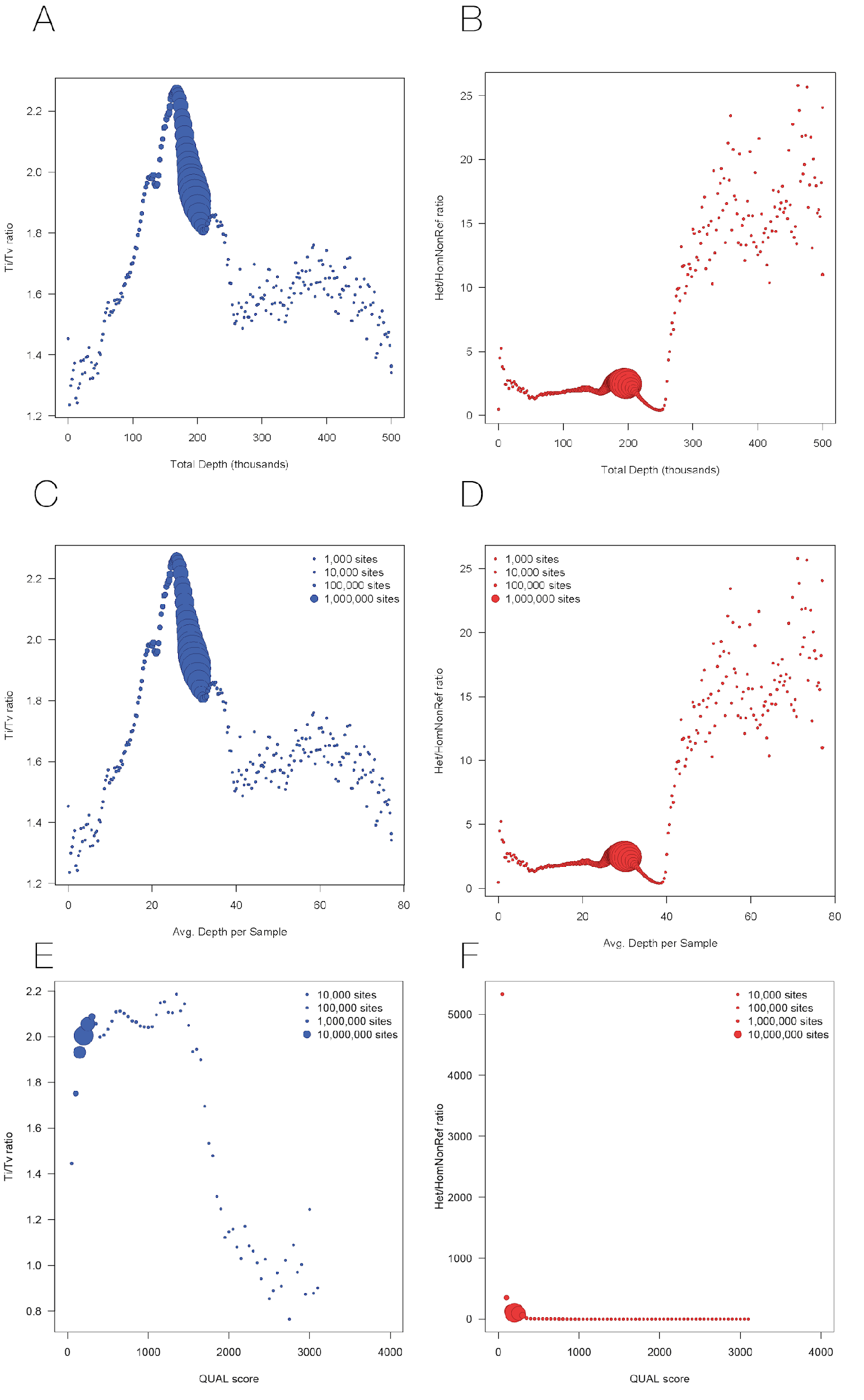

Supplementary Figure 8. SNP-level quality control part 2. Ti/Tv ratio and Het/Hom-non-reference ratios vs Total Depth, Averaged Depth per sample and QUAL score. From these visualizations of the data, we could infer the following QC thresholds: variants with total depth < 10,000 reads (i.e., 1.53X per sample) or > 226,000 reads (i.e., 34.8X/sample) were removed. Variants were not filtered on QUAL score.

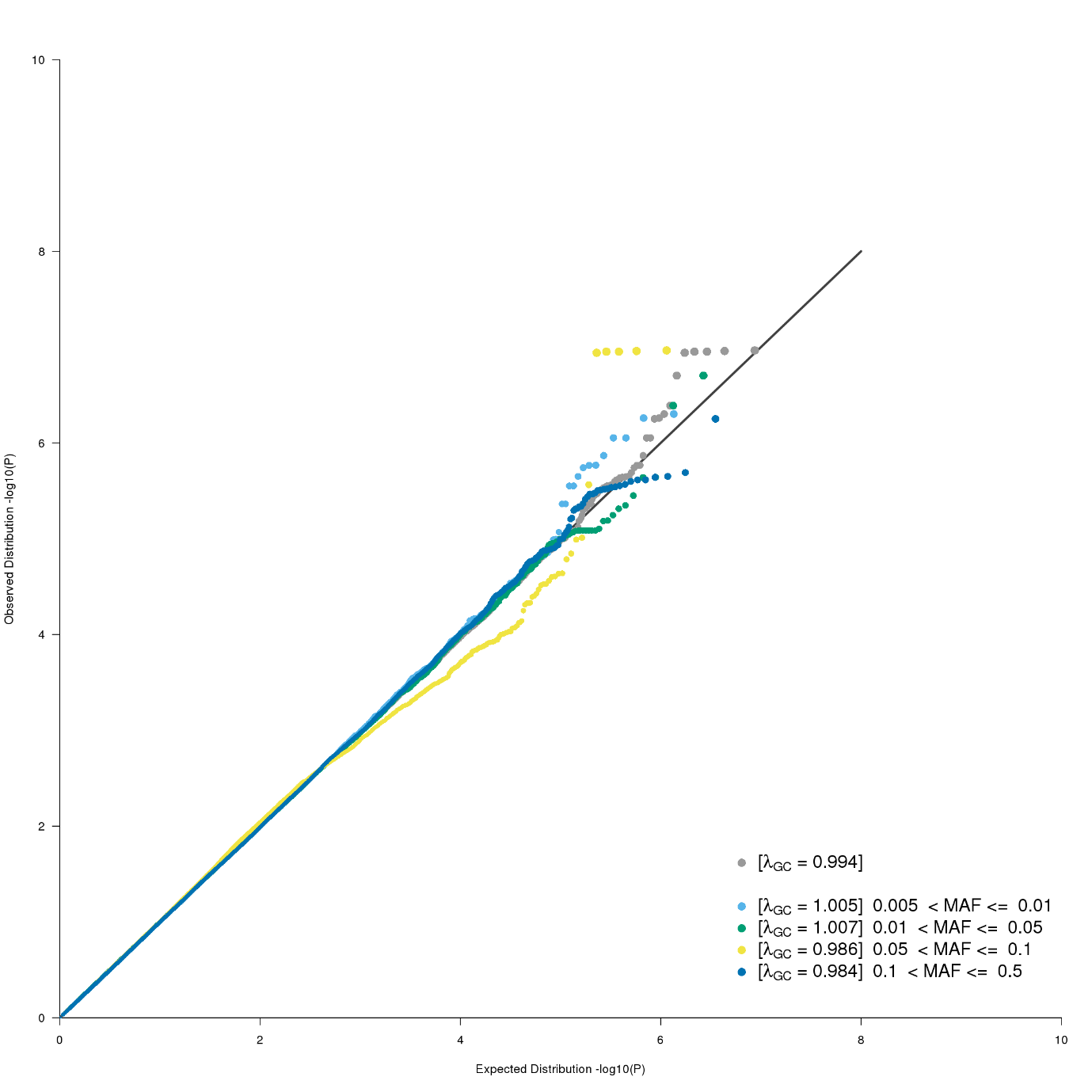

Supplementary Figure 9. SNV QQ plot. QQ plot for single nucleotide variant association using GCTA mlma with a GRM and 20PCs as covariates. Plotted in grey are all SNPs, in colour the same variants stratified by minor allele frequency.

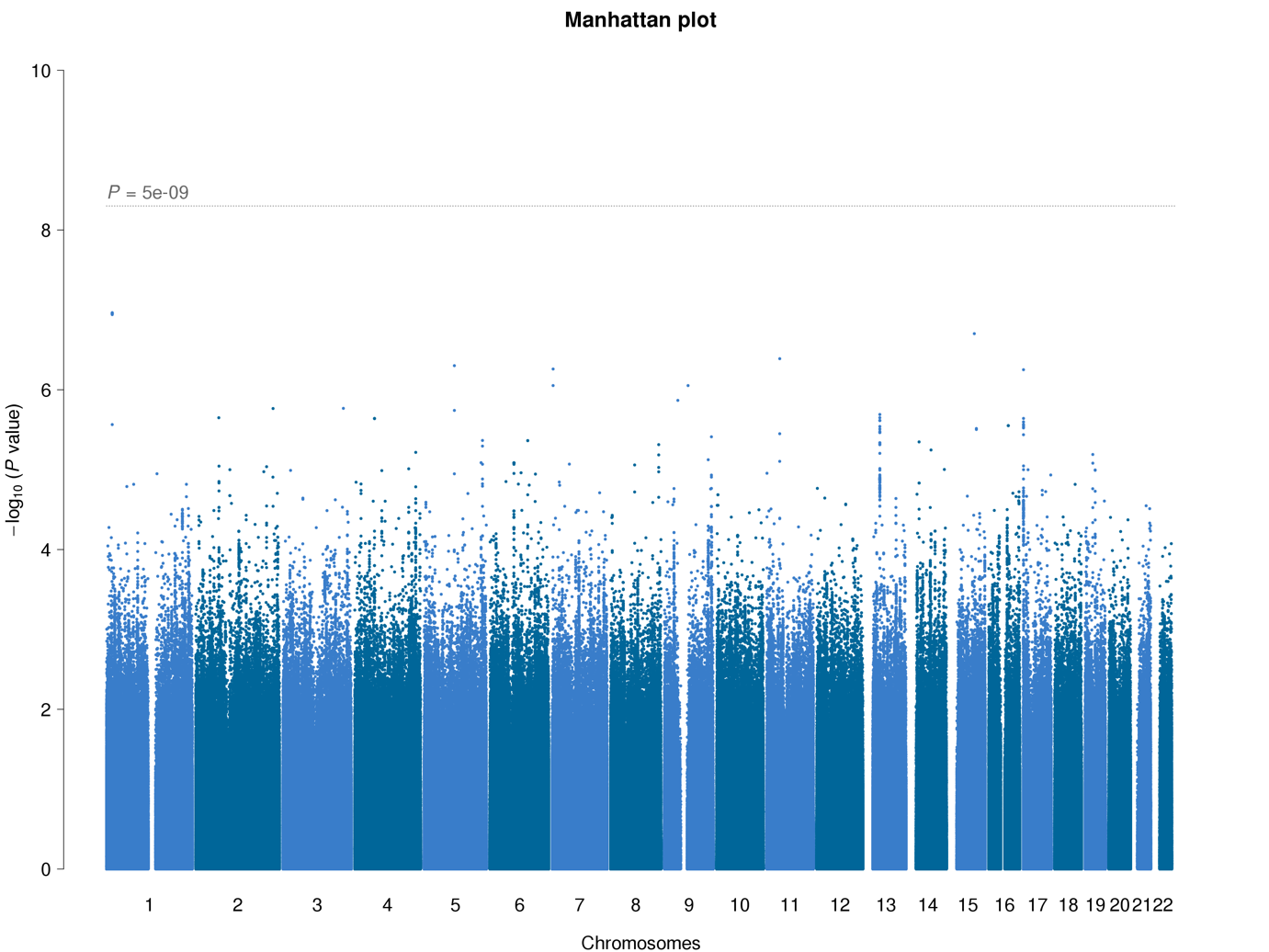

Supplementary Figure 10. SNV associations. Manhattan plot for single nucleotide variant association using GCTA mlma with a GRM and 20PCs as covariates.

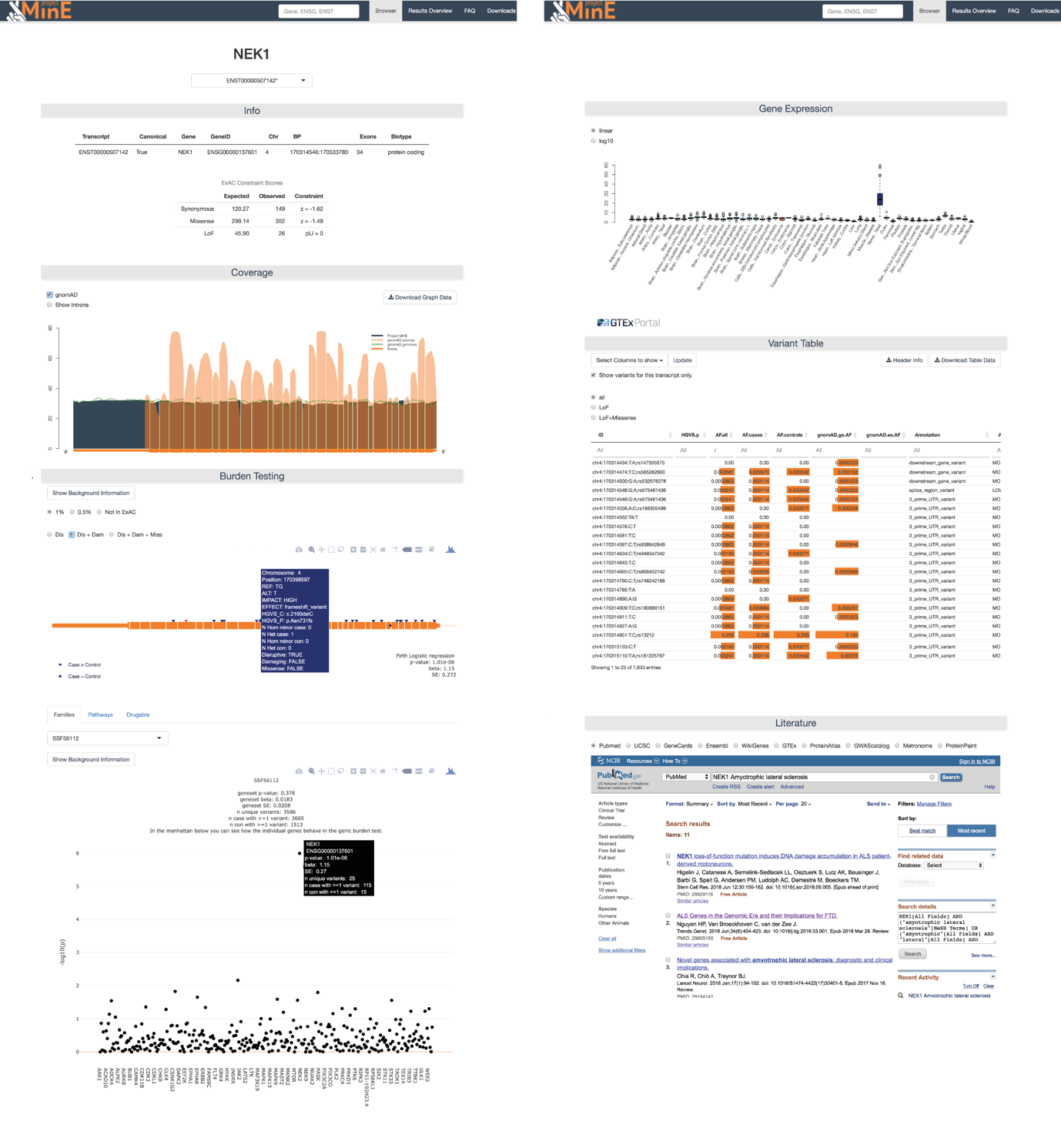

Supplementary Figure 11. Screenshot of the databrowser

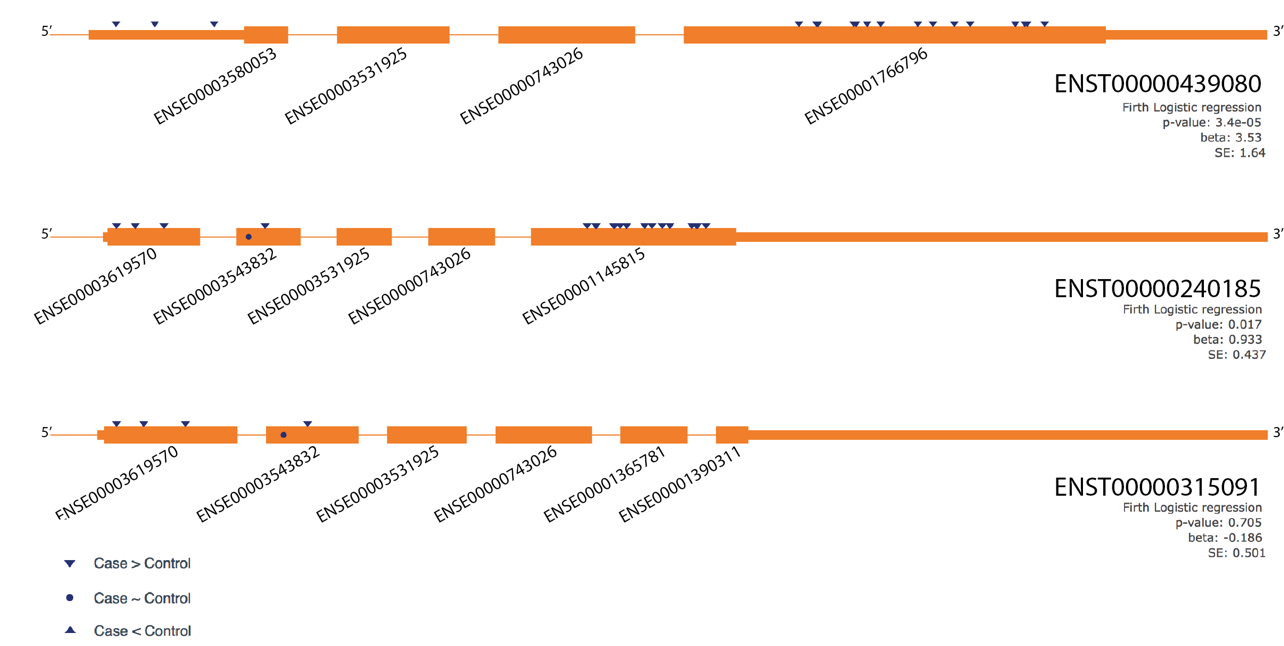

Supplementary Figure 12. Transcript specific burden in TARDBP. All non-synonymous variants (indicated by triangles and circles) with a MAF < 0.5% were included. This overview clearly shows an increased burden in ENSE00001766796 / ENSE00001145815 while the effect might be diluted by exon ENSE00003543832.
